# Interactive Data Integrator for Uniform Data Processing in the MorPhiC Program

**DOI:** 10.64898/2026.10.16.759756

**Authors:** Patrick McKeever, Jesse Flores, Dylan Baker, Ling-Hong Hung, Bryce Nobutoshi Fukuda, Alexandros Orges Koci, Anu Shivalikanjli, Dušica Vidović, Helen Parkinson, Stephan Schürer, Ka Yee Yeung

## Abstract

We present an interactive Data Integrator to facilitate uniform data processing and integration with datasets generated by the NIH funded MorPhiC (Molecular Phenotypes of Null Alleles in Cells) program. In the dataset identification step, users can select published bulk RNA sequencing (RNA-seq) datasets generated by the MorPhiC program, specify external datasets from the Gene Expression Omnibus (GEO) repository, or upload raw RNA-seq data. Subsequently, users can select a subset of samples and indicate experimental design by assigning groups to samples. External data will then be processed using the MorPhiC approved bulk RNA-seq analytical pipeline, and differential expression will be inferred. Users also have the option to download the uniformly processed counts table.

## 1 Introduction

There is a knowledge gap for genes that are underrepresented in the literature. To provide comprehensive functional characterization of human genes, the NIH-funded MorPhiC (Molecular Phenotypes of Null Alleles in Cells) Consortium was initiated in September 2022 to create a comprehensive catalog of the molecular and cellular phenotypes associated with null alleles of all protein-coding human genes (4). This is made possible by recent advances in genetic engineering and cellular models that enable scalable perturbations of gene function in human pluripotent stem cells (iPS cells). The MorPhiC Consortium currently consists of four data production centers (DPCs), one data resource and administrative co-ordinating center (DRACC) and three data analysis and validation centers (DAVs). Each DPC generates high-throughput phenotype data using different null-allele generation strategies across different cellular systems.

Since the initiation of this program, different types of high-throughput data have been generated by the DPCs, including bulk RNA sequencing (RNA-seq), single cell RNA sequencing (scRNA-seq), perturb-seq, ATAC-seq, SLAM-seq, gRNA enrichment and metabolomics data. The DRACC have been developing data processing pipelines to uniformly process data generated by different DPCs, while the DAVs are performing downstream analysis using the processed data. As of September 2026, there have been 4 public data releases from the MorPhiC Data Portal (3; 2; 1).

The National Human Genome Research Institute (NHGRI) has identified MorPhiC as a “community resource project” (4). Therefore, data generated from the consortium are deposited into public repositories, and other resources (such as software, laboratory protocols and cell lines) generated as part of the project are also openly distributed under CC BY 4.0. This paper presents an interactive Data Integrator that facilitates meta-analysis of bulk RNA-seq data generated by the MorPhiC program with data sources from other studies, thus encouraging community driven collaborative data analysis and reuse. Specifically, users can upload their own data or specific external GEO datasets to be processed with the MorPhiC approved analytical pipeline, so as to create a harmonized integrated dataset for downstream analysis. See Figure 1 for an overview of our Data Integrator.

**Figure 1:**
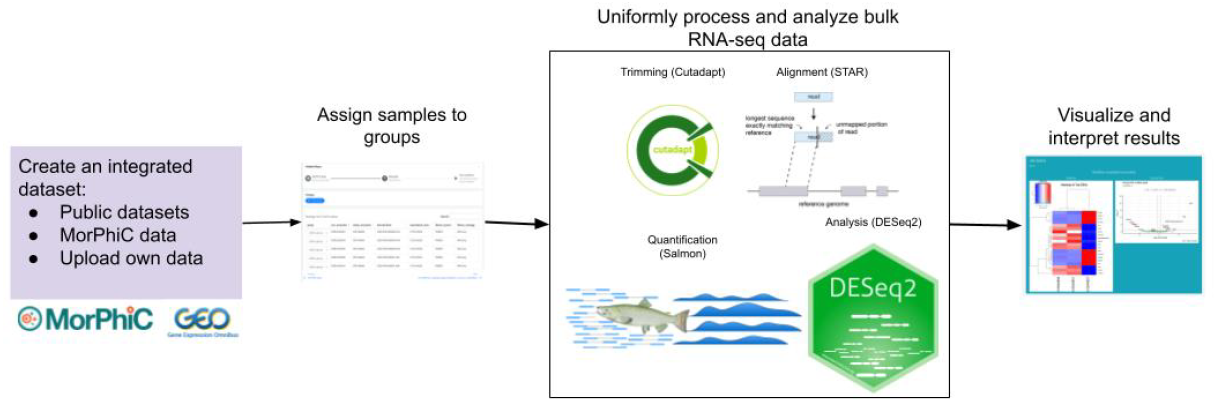
A schematic diagram of our interactive Data Integrator.

## 2 User journey

The user experience begins with the retrieval of studies from the MorPhiC Consortium Public Data Release, as shown in Figure 2. The user can explore and select these published MorPhiC datasets. Moreover, the user can also integrate with non-MorPhiC bulk RNA-seq studies published in GEO by optionally entering GSE accession numbers, as shown in Figure 3. Metadata for the selected datasets is gathered from the NCBI Sequence Read Archive (SRA) and the European Bioinformatics Institute’s European Nucleotide Archive (ENA). The user can define comparison groups, select samples and assign selected samples to groups, as shown in Figure 4. After these samples are selected, users also have the option to uniformly process their own private data by uploading fastq files through the front-end interface, as shown in Figure 5.

**Figure 2:**
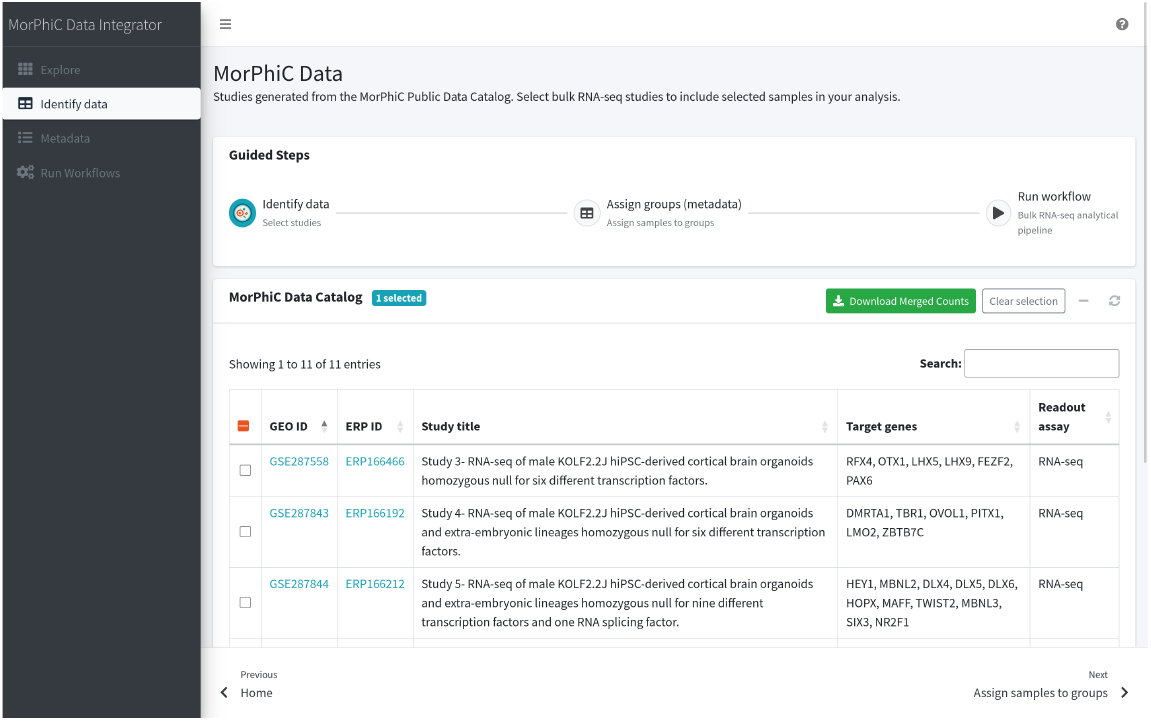
Users can select from the published bulk RNA-seq datasets from MorPhiC.

**Figure 3:**
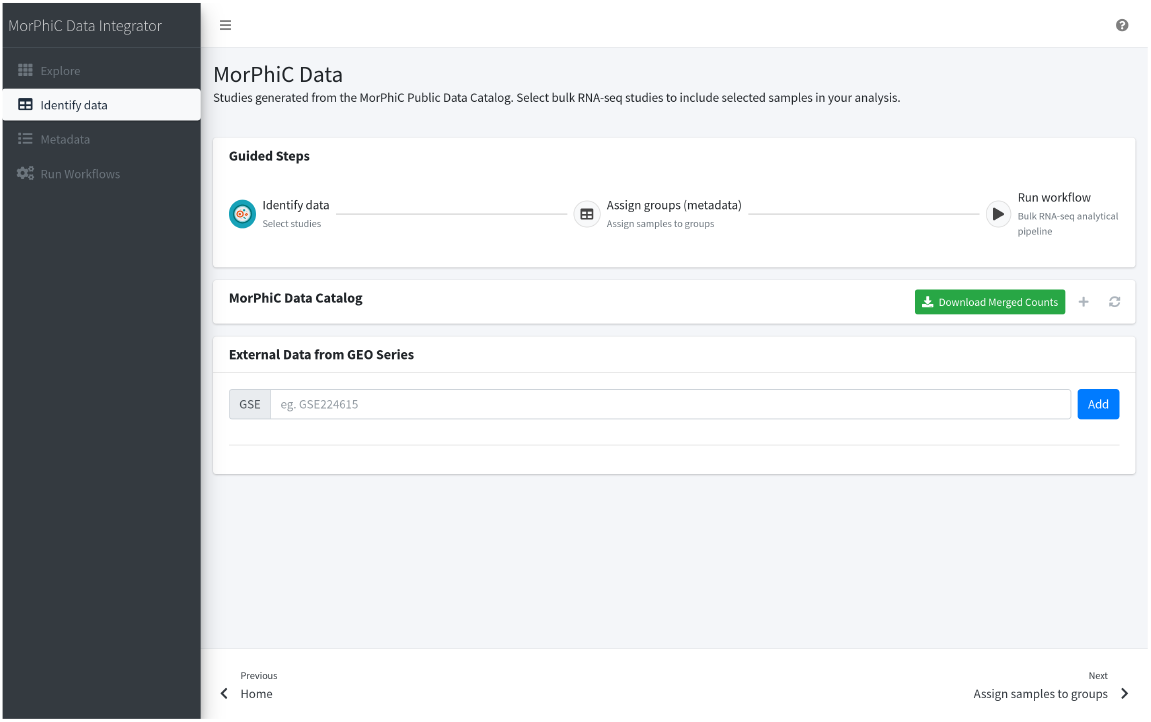
Users can also specify an external dataset from GEO.

**Figure 4:**
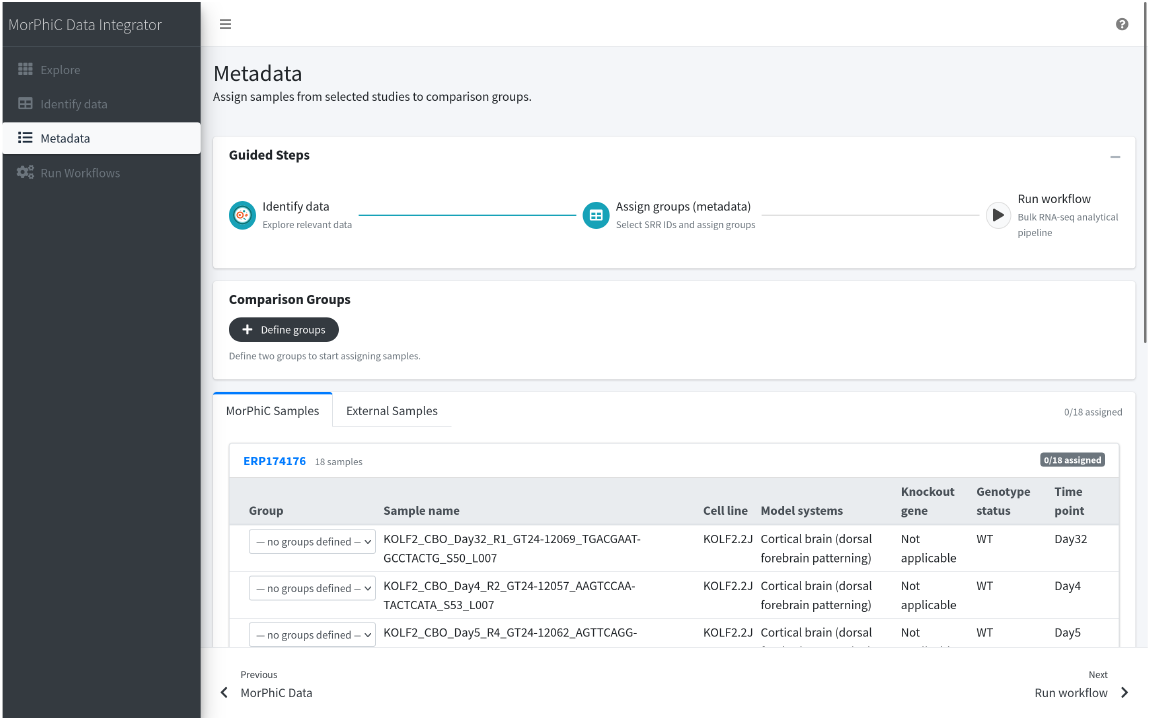
Users can assign groups to selected samples in MorPhiC and external dataset from GEO.

**Figure 5:**
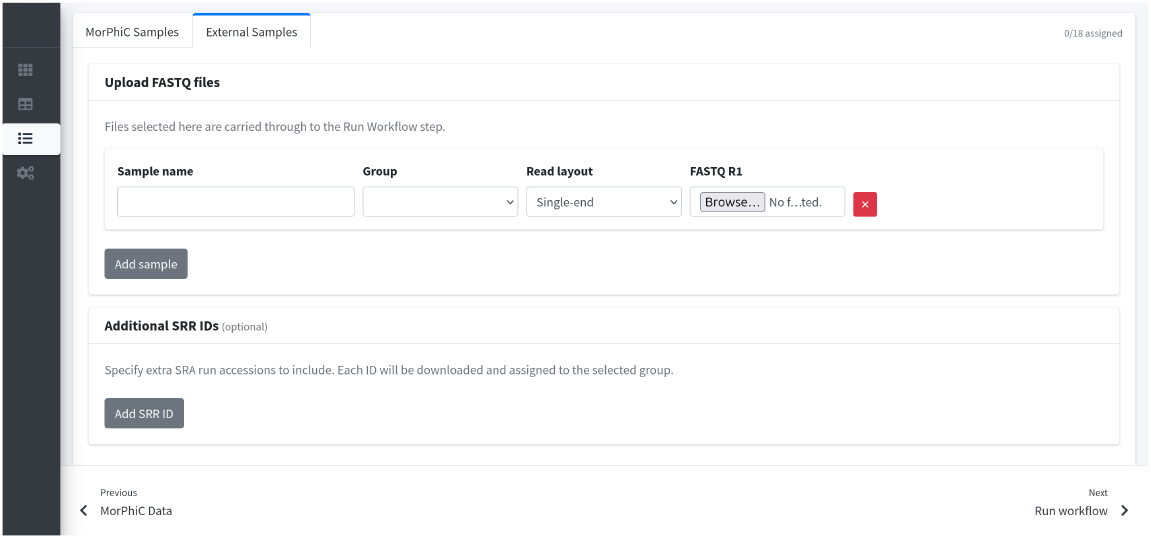
Users can upload and assign groups for private raw sequence data.

Differential expression results are visualized in two complementary ways as shown in Figure 6. A plot is first generated with the EnhancedVolcano package (5), displaying log2 fold change against adjusted p-values with a significance threshold of padj ¡ 0.05, providing an annotated overview of the full results. A heatmap is then produced for the top 20 most significantly differentially expressed genes (ranked by adjusted p-value) using the heatmap.2 (15; 14) function from the gplots package (22), with row-wise z-score scaling applied to the raw count matrix and a blue-red diverging color scheme to indicate relative expression differences across samples.

**Figure 6:**
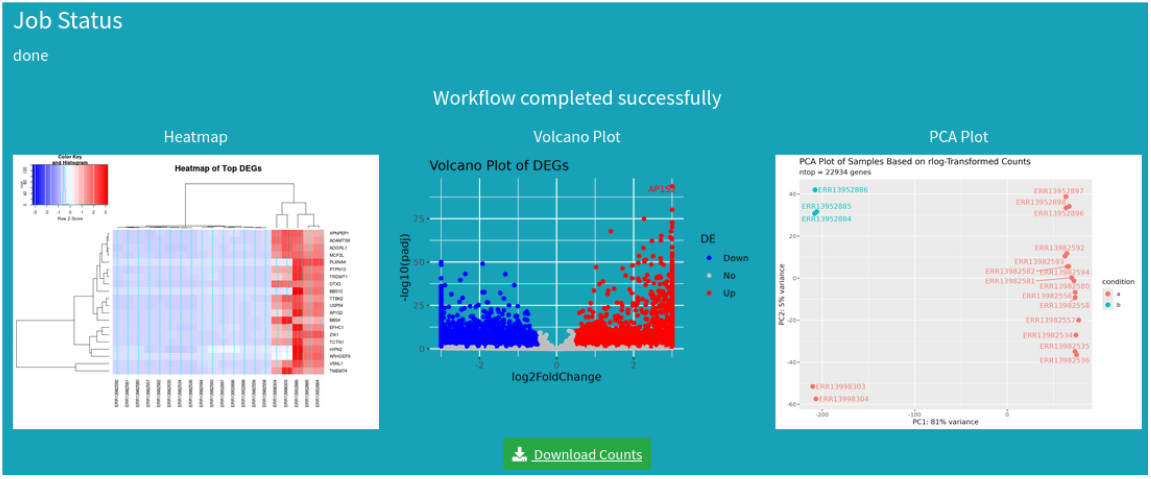
Resulting heatmap and volcano plots from the differential expression analysis.

## 3 Implementation

### 3.1 Graphical user interface

The interactive data integrator utilizes the Flask back-end web framework to create server-side logic and render the front-end. Flask is lightweight and flexible with minimal configuration, allowing for rapid development, and provides support for necessary Python libraries. Once the back-end information retrieval is complete, the front end renders the studies with their information. The user can explore, select studies, enter optional bulk RNA-seq studies published in GEO (GSE accession number), that can then be queued for metadata retrieval. Metadata is gathered from the NCBI Sequence Read Archive (SRA) and the European Bioinformatics Institute’s European Nucleotide Archive (ENA) using the Python package pysradb (6). Each study’s metadata samples are rendered in the front-end, where the user can select and map samples. Once these samples are selected, users can upload one or more fastq files through the front-end interface.

With both the metadata selected and the file uploaded by the user, start scripts are triggered to run a workflow. This process enables the front-end and back-end to exchange status updates which in return present rendered visuals to the user when the workflow is complete.

### 3.2 RNA-seq data processing

Processed counts of user-selected MorPhiC data were retrieved from the European Nucleotide Archive, while the public data from GEO and user-uploaded private bulk RNA-seq data were uniformly processed using a modified version of the MorPhiC Bulk RNA sequencing pipeline run via the Biodepot scheduler (18). The analysis pipeline consists of downloading user-selected SRR IDs via fastqdump, trimming raw FASTQs using Cutadapt (17), aligning to hg38 using STAR (11), and quantifying counts using salmon (20). The workflow has been modified to process a mixture of paired and single-ended FASTQs in case the user-selected samples include both. Following the generation of the counts table, the user has the option to download the uniformly processed counts data integrating MorPhiC and external datasets.

### 3.3 Differential Expression and Visualization

An R script that identifies differentially expressed genes between user-selected conditions is applied to the uniformly processed counts table to generate a heatmap, a volcano plot, and a PCA plot of samples. The analysis pipeline performs differential expression analysis using DESeq2 (16) on the raw count matrices. Sample groupings assigned by the user are used to define the experimental design, with conditions set as the model factor. Ensembl gene IDs are mapped to HGNC gene symbols using biomaRt (13; 12), querying the Ensembl human gene dataset. If no symbol is available, the Ensembl ID is retained as a fallback identifier. Genes are considered differentially expressed at an adjusted p-value threshold of 0.05, with multiple testing correction applied by DESeq2 using the Benjamini-Hochberg procedure.

## 4 Example use cases

### 4.1 Inferring differential expression across MorPhiC datasets

We demonstrated the utility of our Data Integrator to retrieve subsets of processed bulk RNA-seq data across two published MorPhiC datasets. GEO datasets with accession numbers GSE287843 (9) and GSE287844 (10) are bulk RNA-seq data generated by the Jackson Laboratory to investigate the transcriptional effects of knocking out specific regulatory genes in human developmental models. Both datasets utilize the same human induced pluripotent stem cell (hiPSC) line (male KOLF2.2J), the same gene knockout strategy (premature termination codons, PTCs), the same differentiation model systems: cortical Brain Organoid (CBO) and Primitive Syncytium (PrS). GSE287843 (9) consists of a total of 28 samples, among which 10 samples targeting 2 genes (DMRTA1, TBR1) in the CBO model system, and 18 samples targeting 4 genes (OVOL1, PITX1, LMO2, ZBTB7C) in the PrS model system. GSE287844 (10) consists of a total of 48 samples, among which 12 samples targeting 2 genes (SIX3, NR2F1) in the CBO model system, and 36 samples targeting 8 genes (HEY1, HOPX, MAFF, TWIST2, DLX4, DLX5, DLX6, MBNL3) in the PrS model system. Specifically, the 12 samples in the CBO model system consist of 3 null samples per gene and 6 corresponding WT samples and for the PrS there are actually 9 genes (MBLN2 is missing from the list) with 3 null samples each and 9 corresponding WT samples.

In this case study, we selected samples in the CBO model system across 2 MorPhiC datasets (GSE287843 and GSE287844). Specifically, we selected 10 samples from GSE287843 and 12 samples from GSE287844. The 10 samples from GSE287843 consist of 3 null samples per gene (DMRTA1, TBR1) and 4 corresponding WT samples. The 12 samples from GSE287844 consist of 3 null samples per gene (SIX3, NR2F1) and 6 corresponding WT samples. The uniformly processed data can be saved as a CSV text file for further downstream analysis.

### 4.2 Integration with external datasets

We demonstrated the functionality of integrating MorPhiC datasets with external bulk RNA-seq data. In particular, we used an external bulk RNA-seq dataset from the NCBI GEO with accession number GSE99302 that profiles the left and right sides of various brain structures in human foetuses aged 5 to 13 weeks post conception (7; 8; 19). We compared the expression of these fetal and embryonic brain samples (n = 20) with the MorPhiC WT (wild type) brain organoid data from GSE303979 (21) during 6 different time points (at days 3, 4, 5, 20, 25, and 32 post-initiation of differentiation). The resulting PCA is displayed in Figure 7. A clear split among the MorPhiC samples is observable between days 3, 4, and 5 compared to days 20, 25, and 32. Overall, the MorPhiC samples exhibit no clustering with the samples from GSE99302.

**Figure 7:**
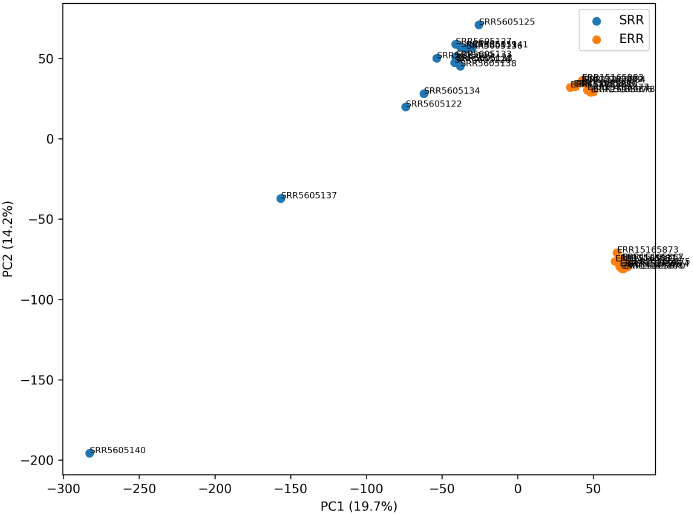
PCA visualization of MorPhiC wild type (red) versus external brain organoid samples from GSE99302 (blue).

## 5 Conclusions

We present an interactive Data Integrator to facilitate uniform data processing and integration with datasets generated by the NIH funded MorPhiC program. Our Data Integrator leverages the Biodepot Scheduler to uniformly process bulk RNA-seq datasets using the consortium approved analytical pipelines. We demonstrated the utility of our tool in two different case studies: across two different MorPhiC datasets and with external bulk RNA-seq datasets. A key limitation of our tool is that only bulk RNA-seq data are considered. Future work includes extending our tool to incorporate additional datasets and analytical pipelines for uniform data processing. Our Data Integrator can also be extended to other contexts, such as integration across data generated by other consortia and programs, and to data from other organisms.

## 6 Data Availability

The data supporting the results of this article are available as follows:

- GSE287843 consists of 28 samples measuring the transcription effects of knocking out 6 genes of the KOLF2.2J line in the CBO and PrS model system. This dataset is available from the MorPhiC Data Catalog at https://morphic.bio/data/ as well as from the NCBI GEO repository with accession number GSE287843 (9).
- GSE287844 consists of 48 samples measuring the transcription effects of knocking out 10 genes of the KOLF2.2J line in the CBO and PrS model system. This dataset is available from the MorPhiC Data Catalog at https://morphic.bio/data/ as well as from the NCBI GEO repository with accession number GSE287844 (10).
- The MorPhiC wild type (WT) bulk RNA-seq dataset profiled male KOLF2.2J hiPSC-derived cortical brain organoids during 6 different time points (at days 3, 4, 5, 20, 25, and 32 post-initiation of differentiation). This dataset is available from the MorPhiC Data Catalog at https://morphic.bio/data/ as well as from the NCBI GEO repository with accession number GSE303979 (21).
- The external bulk RNA-seq dataset is publicly available from the NCBI GEO repository with accession number GSE99302 (19).

## 7 Declarations

### 7.1 List of abbreviations

If abbreviations are used in the text they should be defined in the text at first use, and a list of abbreviations should be provided in alphabetical order.

### 7.2 Ethical Approval

Not applicable.

### 7.3 Competing Interests

L.H.H. and K.Y.Y. have equity interest in Biodepot LLC. The terms of this arrangement have been reviewed and approved by the University of Washington in accordance with its policies governing outside work and financial conflicts of interest in research.

### 7.4 Funding

B.F., J.F., A.O.K., H.P., A.S., D.V., S.S., L.H.H. and K.Y.Y. are supported by National Institutes of Health (NIH) grant U24HG012674. D.M. is supported by NIH grant UM1HG012651. K.Y.Y. is also supported by the Virginia and Prentice Bloedel Endowment at the University of Washington. A.O.K., H.P. and A.S. are also supported in part by EMBL-EBI Core Funds.

### 7.5 Author’s Contributions

P.M. developed the backend for bulk RNA-seq data processing with the scheduler. J.F. designed, implemented and tested the interactive front end. B.F. wrote the differential expression analysis script. L.H.H. designed and developed the MorPhiC approved bulk RNA-seq workflow. P.M. and J.F. prepared the GitHub repository and documentation. D.B. developed the external data integration use case. A.O.K., H.P., and A.S. annotated metadata, brokered bulk RNA-seq datasets to ENA and NCBI GEO. J.F. designed the interactive user interface. D.V. and S.S. contributed to metadata annotations and coordination. K.Y.Y. conceived the study. L.H.H. and K.Y.Y. supervised the study. P.M., J.F. and K.Y.Y. wrote the draft manuscript. All authors read and approved the final manuscript.

## 8 Acknowledgements

We would like to thank Paul Robson and Bill Skarnes at The Jackson Laboratory and NIH grant UM1HG012651 used for generating the MorPhiC datasets. We would like to thank Galabina Yordanova for providing feedback to early versions of the tool.

